# Phage RyR-domain proteins degrade ADPR-based immune signals and fuel NAD^+^ synthesis

**DOI:** 10.64898/2026.05.28.727677

**Authors:** Miguel López Rivera, Renee B. Chang, Colin M. Lewis, Romi Hadary, Julia M. Kovalski, Krista G. Freeman, Zhen-Yu J. Sun, Rotem Sorek, Graham F. Hatfull, Philip J. Kranzusch

## Abstract

Bacterial, plant, and animal cells synthesize nucleotide immune signals as a conserved strategy to defend against viral infection^1–4^. In bacteria, Thoeris anti-phage defense systems convert nicotinamide adenine dinucleotide (NAD^+^) into the cyclic ADP-ribose signals 2′cADPR and 3′cADPR to activate downstream effectors and restrict viral replication^5–8^. Phage proteins can bind and sequester Thoeris signals^6,9–13^, but no mechanisms are known to degrade the exceptionally stable 2′cADPR and 3′cADPR molecules and terminate immune activation. Here we use a forward biochemical screen to discover the mycobacteriophage protein RyDEP as the founding member of an enzyme family that cleaves 2′cADPR and 3′cADPR to inactivate Thoeris defense. We show that RyDEP is a glycosidase that cleaves the ribose-ribose linkage in 2′ and 3′ cADPR immune signals to both inactivate host defense and enable direct restoration of NAD^+^. A crystal structure of the RyDEP–3′cADPR complex in the post-cleavage state explains the molecular basis of immune signal degradation and reveals surprising homology with the Repeat12 domain of animal ryanodine receptors (RyRs) that control calcium flux and muscle contraction^14,15^. We demonstrate that diverse phage RyDEP proteins tune RyR-domain activity to either degrade or sequester immune signals. Our results define RyR-domain proteins as regulators of nucleotide immune signaling and explain how viruses subvert host antiviral defense.

---

Immune proteins across bacterial, plant, and animal evolution respond to viral infection by converting the cellular metabolite NAD^+^ into specialized nucleotide signals that control activation of antiviral effector responses^6–8,16–18^. Toll/interleukin-1 receptor (TIR)-domain containing proteins including *B. cereus* ThsB and *E. coli* ThsB in Thoeris anti-phage defense^6,8^, *N. benthamiana* ROQ1 in plant immunity^7^, *Cr*TIR in algal cells^18^, and SARM1 in human neurons cyclize NAD^+^ by catalyzing formation of ribose-ribose linkages to create 1′′–2′ cyclic ADPR (2′cADPR) and 1′′–3′ cyclic ADPR (3′cADPR) immune signals^17^. Despite the importance of 2′cADPR and 3′cADPR in immune responses across kingdoms of life, no proteins are known to be able to hydrolyze 2′cADPR and 3′cADPR to inactivate signaling. To discover enzymes that disrupt 2′cADPR and 3′cADPR function, we developed a forward biochemical screen of infected-cell lysates to detect viral proteins that degrade ADPR-based signals (Fig. 1a). We synthesized radiolabeled 2’cADPR and 3’cADPR using a two-step enzymatic reaction (Extended Data Fig. 1a–c) and validated each product with comparative liquid chromatography-mass spectrometry (LC-MS) against chemical standards (Extended Data Fig. 1d,e). Next, we incubated labeled 2’cADPR and 3’cADPR with a library of extracts prepared from cells infected with 72 distinct *E. coli, B. subtilis*, and *M. smegmatis* phages and monitored nucleotide immune signal stability using thin-layer chromatography (Extended Data Fig. 2a–c). The resulting lysate-based screen assayed an estimated >10,000 viral proteins and identified putative 2’cADPR cleavage activity in *M. smegmatis* cells infected with phage DyoEdafos (Fig. 1b; Extended Data Fig. 2a–c). Phage DyoEdafos-dependent 2’cADPR cleavage was detected as early as 1.5 h post-infection, consistent with early-to-mid expression of immune evasion proteins observed in other phages (Extended Data Fig. 2d)^19–21^.

**Figure 1.**
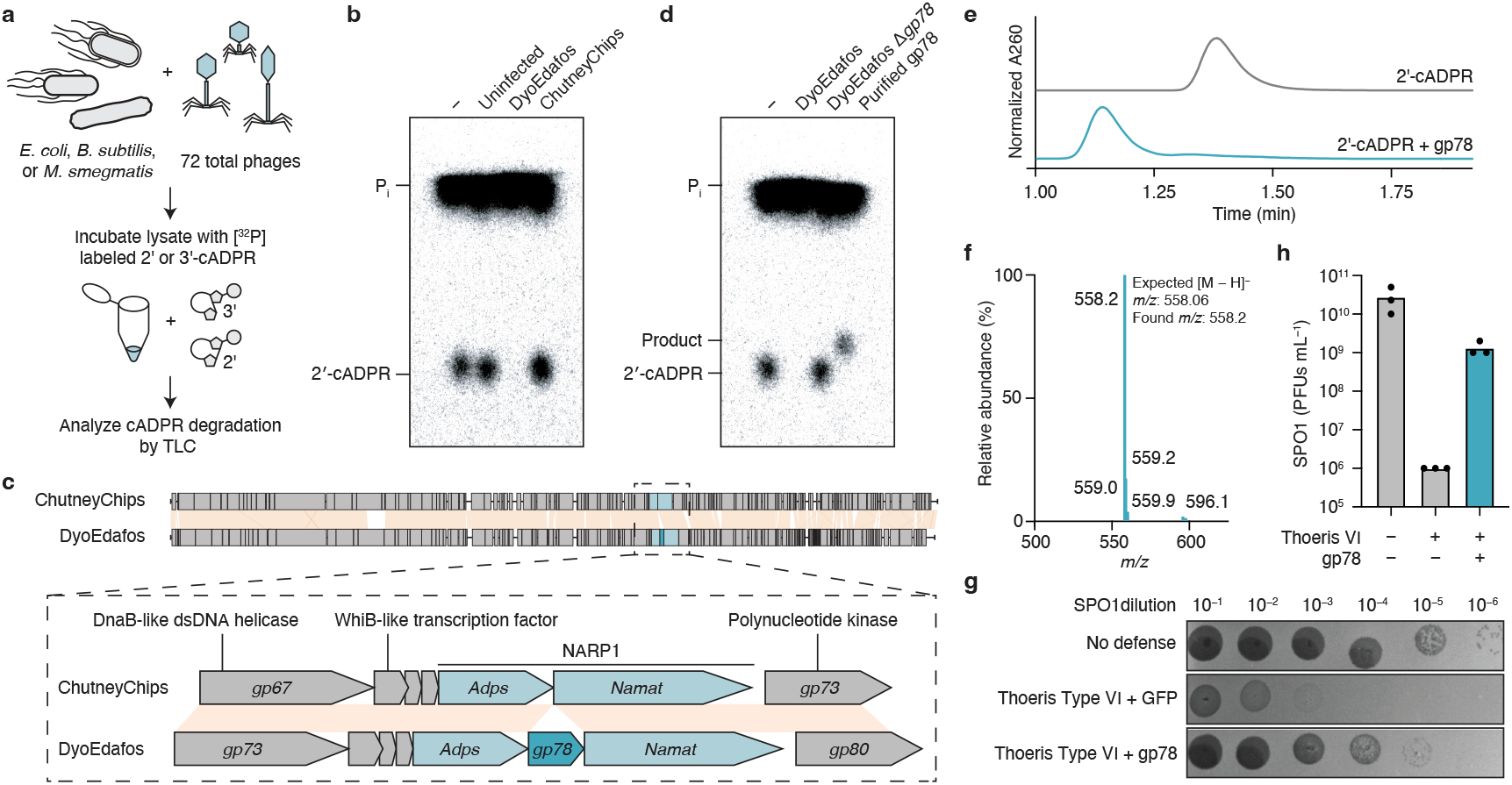
Discovery of RyDEP as a novel anti-Thoeris viral enzyme. **a**, Schematic of biochemical screen to identify cADPR degradation activity in phage-infected lysates using thin-layer chromatography (TLC). **b**, TLC analysis showing cleavage of 2′cADPR by phage DyoEdafos infected lysate, but not by the closely related phage ChutneyChips. Data are representative of two independent experiments. **c**, Comparative genomic analysis of phage DyoEdafos and ChutneyChips genomes identifies gp78 located within the DyoEdafos NARP1 locus as a candidate 2′cADPR-degrading enzyme. Orange shaded regions indicate regions of high nucleotide sequence similarity. **d**, TLC depicting 2′cADPR-processing activity by purified gp78 and lack of activity in phage DyoEdafos Δ*gp78* infected lysates. Data are representative of two independent experiments. **e**, LC-MS trace showing RyDEP-treated 2′cADPR elutes at a different time than standard 2′cADPR. Data are representative of two independent experiments. **f**, LC-MS mass spectra of RyDEP-treated 2′cADPR showing expected m/z values consistent with hydrolyzed 2′cADPR. Data are representative of two independent experiments. **g, h**, Plaque assay (**g**) and quantification (**h**) of *B. subtilis* co-expressing a Type VI Thoeris operon that signals through 2′cADPR with phage DyoEdafos gp78 or a GFP control against serial dilutions of phage SPO1. Data represent three independent experiments.

To identify the phage DyoEdafos protein responsible for 2′cADPR cleavage, we sampled lysates from phylogenetically similar mycobacteriophages. We observed no 2′cADPR cleavage activity in the closely related phage ChutneyChips that is 84% identical to phage DyoEdafos (Fig. 1b), and genome comparison revealed an additional sequence unique to phage DyoEdafos encoding the uncharacterized protein gene product 78 (gp78) (Fig. 1c). Interestingly, bioinformatic analyses of neighboring genes demonstrated that gp78 is inserted within a NARP1 immune evasion operon previously demonstrated in *E. coli* and *Vibrio* phages to encode a two-gene system that synthesizes NAD^+^ to enable evasion of host antiviral NADase effector proteins (Fig. 1c)^22,23^. We engineered a deletion in gp78 and observed that phage DyoEdafos Δ*gp78*-infected cell lysates were no longer capable of degrading 2’cADPR (Fig. 1d). Additionally, we expressed and purified recombinant phage DyoEdafos gp78 protein and confirmed by thin-layer chromatography and LC-MS analysis that this viral protein is alone sufficient to catalyze 2’cADPR degradation (Fig. 1d-f). As tools do not exist in mycobacteria to characterize easily Thoeris defense, we next assessed the role of gp78 in immune evasion in vivo using *B. subtilis* phages known to be susceptible to a type VI Thoeris system that signals through 2’cADPR^8^. Replicative fitness of the Bacillus phages SPO1 and SBSphiJ was restored upon co-expression of phage DyoEdafos gp78 confirming that this protein enables anti-Thoeris immune evasion (Fig. 1g,h, Extended Data Fig. 2e,f). Given these results, and shared structural homology between gp78 and the Repeat12 (R12) RyR-domain of ryanodine receptors (RyRs) (see below), we named this viral protein RyDEP (RyR-domain Defense Evasion Protein). Together, these results demonstrate that RyDEP is a viral enzyme that can degrade cADPR-based signals to evade host antiviral immunity.

## RyDEP glycosidase activity fuels viral NAD^+^ synthesis

To define the mechanism of RyDEP nucleotide immune signal degradation, we analyzed reaction cleavage products in vitro using LC-MS and nuclear magnetic resonance (NMR). RyDEP catalyzed the conversion of 2′cADPR and 3′cADPR into identical cleavage products with the same LC-MS retention time and an m/z value consistent with linearized ADPR (Fig. 2a,b). Enzyme titration experiments demonstrated that RyDEP more efficiently processes 2′cADPR than 3′cADPR, likely explaining why only 2′cADPR cleavage activity was observed in initial phage DyoEdafos lysates experiments (Extended Data Fig. 3a). Cleavage of 2′cADPR and 3′cADPR into identical products suggested that RyDEP degrades the 1′′–2′ or 1′′–3′ ribose-ribose O-glycosidic bond in host nucleotide immune signals. Consistent with this hypothesis, we observed in NMR analyses of cleaved 2′cADPR loss of characteristic J-coupling peaks in the HMBC spectrum (Fig. 2c & Extended Data Fig. 3c)^6^. Additionally, analysis of the NMR COSY spectrum revealed creation of hydroxyl peaks strongly linked to the product 2′ proton (Fig. 2c), demonstrating that RyDEP is a glycosidase that hydrolyzes the ribose-ribose linkage in 2′cADPR and 3′cADPR to generate linear ADPR. Finally, we confirmed using in vivo phage challenge assays that RyDEP enables immune evasion of 3′cADPR signaling in a type I Thoeris system (Extended Data Fig. 3d,e), demonstrating that RyDEP glycosidase activity is sufficient to target both 2′cADPR and 3′cADPR.

**Figure 2.**
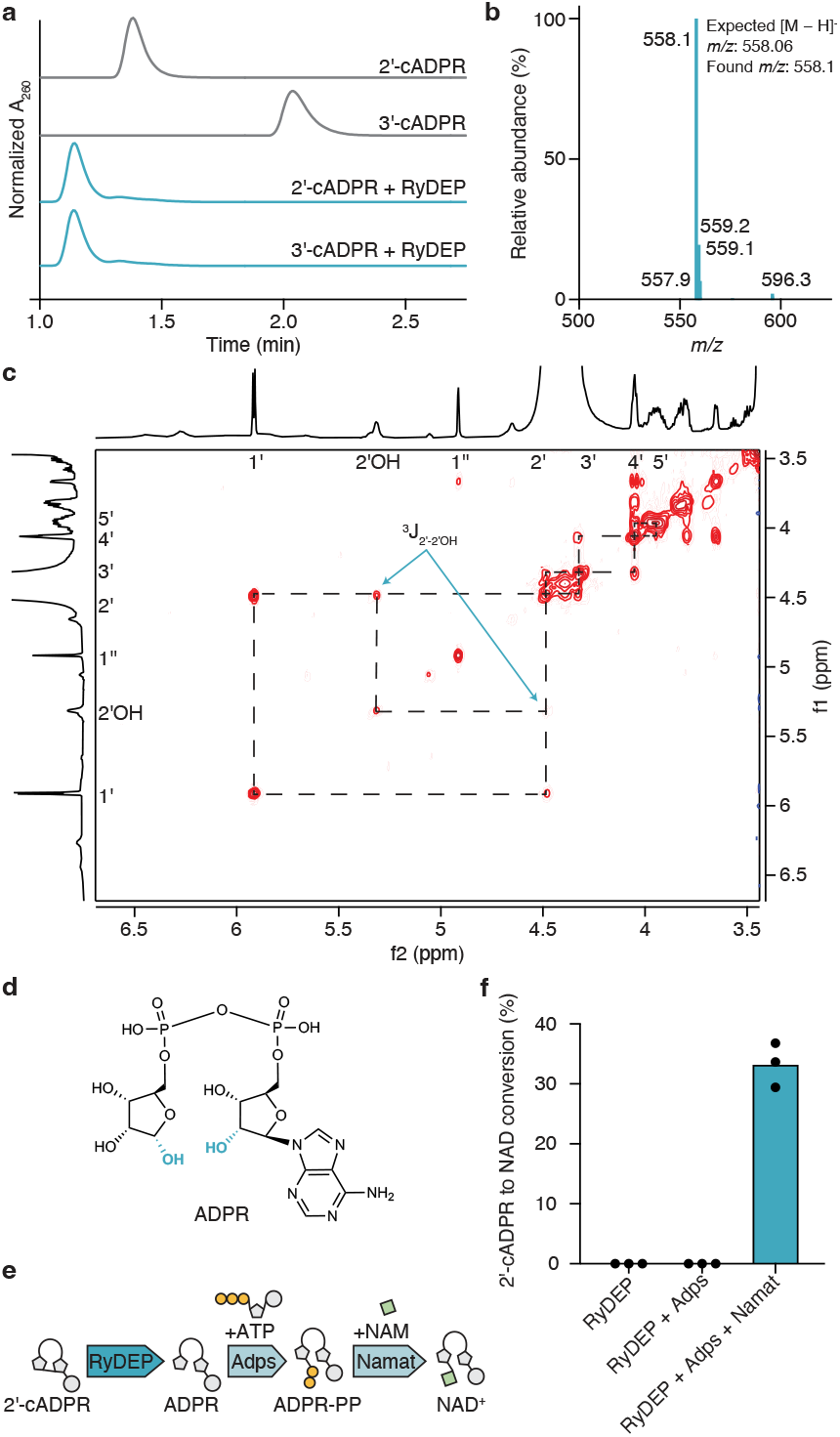
RyDEP is a glycosidase that generates ADPR to fuel NAD^+^ synthesis. **a**, LC-MS trace showing RyDEP-treated 2′cADPR and 3′cADPR elute at the same retention time as a product that is distinct from a 3′cADPR standard. Data are representative of two independent experiments. **b**, LC-MS mass spectra of RyDEP-treated 3′cADPR showing expected m/z values consistent with hydrolyzed 3′cADPR. Data are representative of two independent experiments. **c**, NMR COSY spectrum showing a new hydroxyl peak (washed out by adding a drop of D2O) 3J-coupled to the 2′ proton, consistent with linear ADPR. **d**, Molecular structure of the identified product, linear ADPR, with the hydroxyl groups generated by hydrolysis of the O-glycosidic bond highlighted in blue. **e**, Schematic of the NARP1 pathway synergizing with RyDEP to convert 2′cADPR into NAD^+^. ADPR-PP stands for ADP-ribose pyrophosphate, NAM stands for nicotinamide. **f**, Conversion of 2′cADPR into NAD^+^ by purified phage DyoEdafos NARP1 enzymes and RyDEP. NAD^+^ concentrations were determined using the NAD/NADH-Glo assay. Data are representative of three independent experiments.

We next investigated potential synergy between RyDEP and the activity of neighboring phage DyoEdafos NARP1 operon proteins in viral immune evasion. The phage NARP1 enzymes Adps and Namat were previously shown to catalyze NAD^+^ synthesis from linear ADPR produced by host NADase effector proteins^22^. The RyDEP 2′cADPR and 3′cADPR cleavage product is identical to the precursor linear ADPR metabolite required for input into NARP1 operon activity. We therefore reconstituted the phage pathway in vitro and observed that co-incubation of RyDEP-treated 2′cADPR with recombinant phage DyoEdafos Adps and Namat proteins allowed conversion of 2′cADPR into NAD^+^ (Fig. 2f). Together, these data show that RyDEP is a glycosidase that hydrolyzes the ribose-ribose O-glycosidic bond in cADPR-based host immune signals to generate linear ADPR and that this product can be directly used by partnering NARP1 enzymes to reconstitute NAD^+^ synthesis and restore viral replication.

## RyDEP shares homology with a domain in animal RyRs

We next determined a 1.7 Å crystal structure of phage DyoEdafos RyDEP in complex with 3′cADPR (Extended Data Table 1). The RyDEP–3′cADPR structure represents a post-reaction state and reveals a homodimer with two opposing castanet-like protomers, each harboring a molecule of linear ADPR within a deep central pocket (Fig. 3a). The RyDEP dimeric interface is formed by helices α1 and α4 that interlock and bury 1,550 Å2 surface area (Fig. 3a). RyDEP helices α2 and α3 project away from the dimeric interface to create the surface-exposed pocket for nucleotide immune signal recognition (Fig. 3a). We used DALI analysis to compare RyDEP to structures in the Protein Data Bank and discovered strong homology with the R12 regulatory domain of animal RyR proteins including human RyR3 (Z-score 8) and a putative ryanodine receptor-like protein in *B. thetaiotaomicron* (Z-score 6.1)^24^. Human RyR3 is a calcium channel in the endoplasmic reticulum with key roles in skeletal muscle cells in early development and adult neuronal cell function^25–29^. The RyR region sharing structural homology with RyDEP is a similar two-fold symmetric globular domain formed by tandem repeats in the channel cytosolic periphery (Extended Data Fig. 4a)^25–27^. Comparison of RyDEP and a previous structure of human RyR3 in complex with adenine-containing ligands and the small molecule inhibitor dantrolene further reveals a shared mechanism of ligand recognition^15^. In both RyDEP and the RyR3 R12 domain, ligand recognition occurs in a cavity formed by four central helices and a loop braced between helices α1–α2, with an additional loop involved in ligand recognition between helices α3–α4 in RyDEP (Fig. 3c and Extended Data Fig. 4b,c). In the RyDEP structure, 3′cADPR ligand recognition occurs through a series of interactions including a histidine residue H75 and an aspartic acid residue D117 that contact the terminal ribose of the ligand, a tryptophan residue W78 that braces the face of the adenine base, and an additional conserved histidine residue H100 that coordinates the ligand phosphates (Fig. 3d,e). Each of these ligand-binding residues are conserved in animal RyR R12 domain proteins and the corresponding residues in human RyR3 form remarkably similar ligand-recognition interactions with dantrolene and adenine ligands (Fig. 3d,e). Broader phylogenetic analysis demonstrates conservation of substrate binding residues across the R12-domain protein family (Pfam PF02026) with RyDEP clustering within a clade of sequences predominantly of bacterial and viral origin (Extended Data Fig. 4b,c). Together, these data identify RyDEP as a glycosidase enzyme member of the R12-domain protein family and demonstrate an evolutionary link between viral immune evasion and conserved nucleotide-recognition modules found across domains of life.

**Figure 3.**
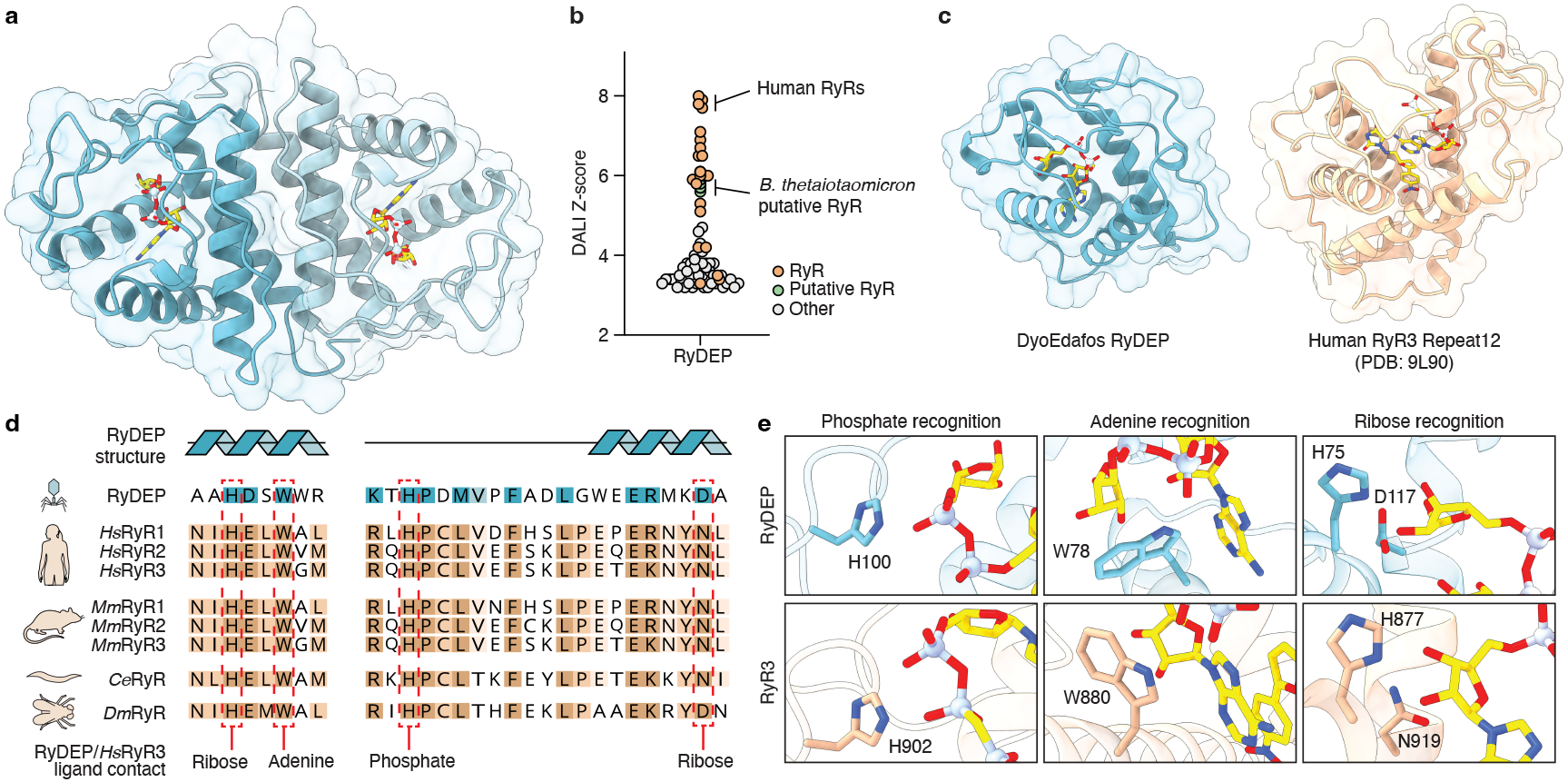
Homology to RyR domain reveals conserved mechanism of nucleotide recognition. **a**, Overview of RyDEP dimer from phage DyoEdafos in complex with 3′cADPR in the post-cleavage state. **b**, DALI Z-scores for the top hits for the phage DyoEdafos RyDEP protomer when searched against all entries in the Protein Data Bank. **c**, Comparison of the phage DyoEdafos RyDEP protomer in complex with 3′cADPR in the post-cleavage state (blue) with a human RyR3 R12 domain crystal structure in complex with dantrolene and the nonhydrolyzable ATP analog AMP-PCP (PDB 9L90, orange). **d**, Sequence alignment of RyDEP with the first repeat in the R12 domain of human (*Hs*), mouse (*Mm*), *C. elegans* (*Ce*), and *D. melanogaster* (*Dm*) RyRs. Highlighted in red are key conserved residues involved in recognizing adenine ligands in both DyoEdafos RyDEP and human RyR3 R12. The RyDEP secondary structure is mapped above the alignment, with helices shown as blue cartoons. **e**, Close-up views of conserved residues in DyoEdafos RyDEP and human RyR3 R12 that mediate recognition of adenine-containing ligands in a similar manner.

## Catalysis and sequestration define Thoeris evasion by phage RyR-domain proteins

To define the mechanism of RyDEP 2′cADPR and 3′cADPR hydrolysis, we examined the ADPR product and surrounding active-site residues in the post-reaction structure. Clear electron density allowed unambiguous assignment of an inverted configuration at C1 of the terminal ribose of ADPR, a hallmark feature of inverting glycosidase enzymes (Fig. 4a, Extended Data Fig. 5a)^30,31^. Analysis of the RyDEP cleavage product NMR spectra further confirmed the inverted α-anomer conformation of the ADPR terminal ribose compared to the β-anomer of the ADPR adenosine ribose (Extended Data Fig. 3c). Consistent with the acid-base chemistry of glycosidase enzymes^30–32^, the RyDEP active site is composed of a catalytic dyad formed by residue H75 positioned to activate a water molecule for backside nucleophilic attack of the ribose C1 and residue D117 positioned to protonate the leaving group to hydrolyze 2′cADPR or 3′cADPR into linear ADPR (Fig. 4b). We mutated conserved amino acids in the RyDEP active site and confirmed the catalytic dyad and surrounding ligand-coordinating residues are required for 2′cADPR degradation in vitro, and that mutation of the catalytic H75 residue abolishes RyDEP-mediated evasion of Thoeris defense in vivo (Fig. 4c,d, Extended Data Fig. 5b–e).

**Figure 4.**
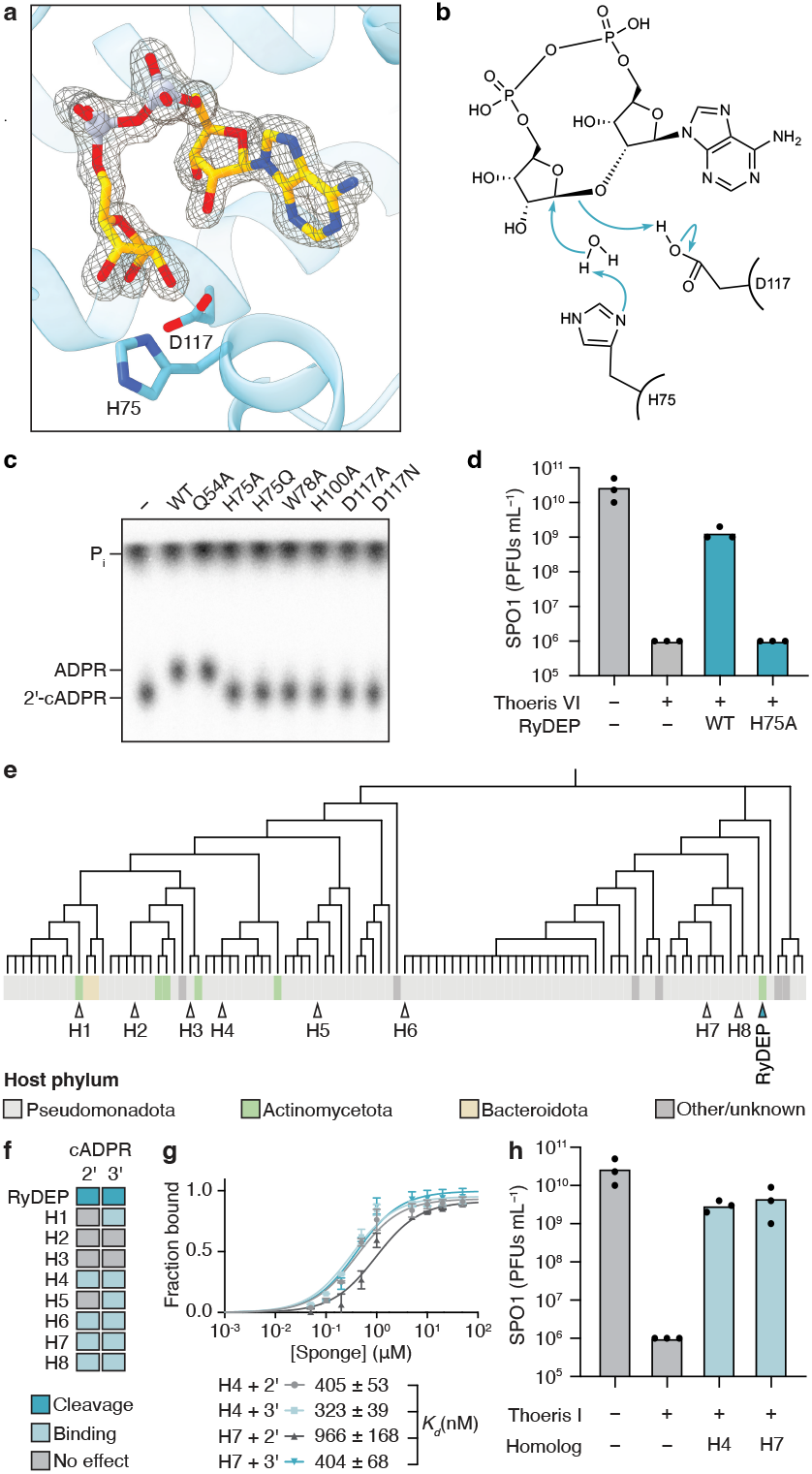
RyDEP catalysis and signal sequestration by homologs define Thoeris evasion. **a**, Electron density map of the ADPR product bound in the RyDEP active site following 3′cADPR cleavage, with the catalytic residues H75 and D117 shown as sticks. **b**, Proposed inverting glycosidase mechanism for RyDEP-catalyzed hydrolysis of the 2′cADPR O-glycosidic bond. **c**, TLC analysis of 2′cADPR hydrolysis by RyDEP active-site point mutants. Data are representative of two independent experiments. **d**, Plaque assay quantification of phage SPO1 replication in B. subtilis coexpressing Type VI Thoeris with wild-type RyDEP or the catalytically inactive H75A mutant. Data represent three independent experiments. **e**, Phylogenetic tree of virally encoded R12-domain proteins in the RyDEP family (Pfam PF02026). Host phyla are indicated by colored rectangles below each tree node. Eight homologs selected for biochemical characterization (H1–H8) are indicated with open triangles, and DyoEdafos RyDEP is indicated with a filled blue triangle. **f**, Summary of EMSA binding analysis showing the ability of each RyDEP homolog to form complexes with 2′cADPR and 3′cADPR. Summarized data represent two independent experiments. **g**, EMSA quantification of H4 and H7 binding to 2′cADPR and 3′cADPR. Fraction bound was calculated as bound signal divided by total signal at each ligand concentration and fit to a one-site binding model. Data represent two independent experiments. **h**, Plaque assay quantification of phage SPO1 replication in B. subtilis co-expressing Type I Thoeris with the selected RyDEP homologs H4 and H7 or a GFP control. Data represent three independent experiments

We next constructed a phylogenetic tree of all virally encoded R12-domain proteins in the RyDEP family (Pfam PF02026) and investigated the functional diversity of RyDEP homologs across distinct phages (Fig. 4e). We selected eight RyDEP homologs representing each major branch of the protein tree, purified the R12-like domains, and tested the ability of these proteins to bind and degrade 2′cADPR and 3′cADPR signals. Six of eight tested RyDEP homologs formed a stable complex with 2′cADPR/3′cADPR signals in an electrophoretic mobility shift assay (Fig. 4f, Extended Data Fig. 6a,b). However, despite high sequence conservation with phage DyoEdafos RyDEP, the other tested RyDEP homologs did not catalyze 2′cADPR and 3′cADPR degradation, suggesting these proteins may not rely on 2′cADPR and 3′cADPR hydrolysis to inactivate Thoeris defense (Fig. 4f, Extended Data Fig. 6a,b). We characterized homolog 4 from a *Ruegeria* phage and homolog 7 from a *Pectobacterium* phage, and observed that both RyDEP proteins bound 2′cADPR and 3′cADPR with nanomolar dissociation constants similar to other characterized phage immune evasion proteins named sponges that tightly bind and sequester nucleotide immune signals to inactivate host defense (Fig. 4g, Extended Data Fig. 6c,d)^9,10,12,33–35^. In contrast to the glycosidase-type RyDEP homolog from phage DyoEdafos that more efficiently targets 2′cADPR, sponge-type RyDEP homologs exhibited higher affinity for 3′cADPR than 2′cADPR (Fig. 4g, Extended Data Fig. 6c,d). Using phage challenge assays, we observed that both sponge-type RyDEP homologs were sufficient to enable immune evasion in vivo and exhibited enhanced activity against 3′cADPR-signaling in type I Thoeris anti-phage defense (Fig. 4h and Extended Data Fig. 6e-g), supporting that phages can tune the biochemical properties of a single family of immune evasion proteins to either sequester or degrade host nucleotide immune signals.

Our study reveals RyDEP as the founding member of a family of virally encoded RyR-domain proteins that degrade or sequester cADPR-based signals to evade Thoeris antiphage defense. Phage DyoEdafos RyDEP selectively hydrolyzes the ribose–ribose glycosidic bond in 2′cADPR and 3′cADPR, establishing that viruses can use glycosidases as an additional enzyme class to inhibit host nucleotide immune signals^12,19,36–40^. We demonstrate that RyDEP glycosidase activity hydrolyzes 2′cADPR and 3′cADPR into linear ADPR and can synergize with NARP1 operon enzymes to further regenerate intact NAD^+^. In this way, RyDEP activity allows phages to convert a host immune signal that inhibits viral replication into a metabolite essential for productive infection. The ability of a virus to restore cellular metabolites manipulated in host defense could be particularly advantageous in slow-growing hosts such as mycobacteria, where limited cytosolic NAD^+^ availability may constrain phage replication^41,42^. Investigation of RyDEP homologs reveals members of this protein family can also function as sponge proteins that bind and sequester host immune signals. Functional variation of RyDEP homologs as either hydrolase enzymes or sponge proteins suggests that other large families of immune evasion sponge proteins that target nucleotide immune signals may have enzymatic members^9,10,33–35^. Finally, unexpected structural and sequence homology between phage RyDEP and the R12-domain of animal RyR ion channels suggests additional features of nucleotide immune signal regulation potentially shared across kingdoms of life. cADPR has previously been implicated in modulation of animal RyR protein function^25–29^, and the structural homology between RyDEP and RyR R12 domains now supports future studies on potential interactions between this region of RyR receptors and cADPR-based signals. Together, our results define RyDEP as an immune evasion glycosidase that inhibits Thoeris anti-phage defense and establish viral members of the R12-domain protein family as factors that regulate nucleotide immune signaling.

## Acknowledgements

The authors are grateful to members of the Kranzusch lab for helpful comments and discussion. We thank the students and faculty at the University of Houston Downtown within the SEA-PHAGES program for the isolation and characterization of DyoEdafos. We thank D. Wassarman, S. Yamaguchi, and Y. Li from the Kranzusch laboratory for assistance with LC-MS analysis, and S. Fernandez from the Kranzusch laboratory for assistance with phage genome sequencing. The work was funded by grants to P.J.K. from the Pew Biomedical Scholars program, the Burroughs Wellcome Fund PATH program, The G. Harold and Leila Y. Mathers Charitable Foundation, The Mark Foundation for Cancer Research, the Cancer Research Institute, the Parker Institute for Cancer Immunotherapy, the Gates Foundation (INV-083469), and the National Institutes of Health (1DP2GM146250-01), grants to R.S. from the European Research Council (ERC-AdG GA 101018520), the Israel Science Foundation (MAPATS Grant 2720/22), the Deutsche Forschungsgemeinschaft (SPP 2330, grant 464312965), and a research grant from Magnus Konow in honor of his mother Olga Konow Rappaport, grants to G.F.H. from the National Institutes of Health (GM131729) and the Howard Hughes Medical Institute (GT12053). M.L.R. is supported through the Rafael del Pino Excellence Scholarship Programme and a Real Colegio Complutense at Harvard University Scholarship. R.B.C. is supported through a Landry Cancer Biology Research Fellowship (Harvard Faculty of Arts and Sciences). X-ray data were collected through support by an agreement between the Advanced Photon Source, a US Department of Energy (DOE) Office of Science user facility operated for the DOE Office of Science by Argonne National Laboratory under contract no. DE-AC02-06CH11357, through the Northeastern Collaborative Access Team beamlines, which are funded by the National Institute of General Medical Sciences from the National Institutes of Health (P30 GM124165) and a NIHORIP HEI grant (S10OD021527). NMR work was performed at Harvard Medical School Bio-molecular NMR facility and Dana-Farber Cancer Institute NMR core.

## Author contributions

The study was designed and conceived by M.L.R. and P.J.K. RyDEP discovery, crystallography, and biochemical experiments were performed by M.L.R. with assistance from R.B.C. Phage infection and genome engineering experiments were performed by C.M.L., J.M.K., K.G.F., G.F.H., and M.L.R. Phage Thoeris immune evasion assays were performed by R.H. and R.S. Z.-Y.J.S. performed NMR analysis. The manuscript was written by M.L.R. and P.J.K. All authors contributed to editing the manuscript and support the conclusions.

## Data availability statement

Coordinates and structure factors of the RyDEP–3′cADPR complex have been deposited in PDB under the accession code 35TM. All other data are available in the manuscript or the supplementary information.

## Materials and methods

### Bacterial and phage strains

For the phage defense assays, Type I Thoeris from *Bacillus cereus* MSX-D12 and type VI Thoeris from the metagenomic *Bacillus* scaffold (TIR ID: Ga0102589_100169733), were previously cloned under their native promoters into the *amyE* locus of the *B. subtilis* BEST7003 genome^5,8^. A GFP-expressing vector was integrated as a negative control in the *thrC* locus. RyDEP and RyDEP homologs and mutants were cloned under the control of an IPTG-inducible promoter into the thrC-Phspank vector^6^, which carries a low-copy pSC101 origin of replication and a kanamycin resistance marker^12,13,43^. The DNA sequences of RyR proteins RyDEP, RyDEP H75A, H4 (*Ruegeria* phage RyDEP homolog) and H7 (*Pectobacterium* phage RyDEP homolog) were PCR-amplified from constructs synthesized by Twist Biosciences (see below). Plasmids were transformed into *E. coli* DH5α with the custom protocol and extracted with Qiagen miniprep kits. The plasmids were sequenced with Plasmidsaurus and introduced into *B. subtilis* BEST7003 cells in the *thrC* locus as previously described^6^, where the respective defense system was integrated into the *amyE* locus. Transformations were confirmed by PCR.

The *B. subtilis* phage SBSphiJ (GenBank: LT960608.1) was previously isolated by the Sorek lab^5^ and SPO1 (BGSC: 1P4; GenBank: NC_011421.1) was obtained from the BGSC. All *B. subtilis* phages were propagated on *B. subtilis* BEST7003 by picking a single phage plaque into a liquid culture grown at 37 °C to an optical density at 600 nm (OD600) of 0.3 in MMB broth until culture collapse (or 3 h in the case of no lysis). The culture was then centrifuged for 10 min at 3,200 × g and the supernatant was filtered through a 0.2 µm filter to get rid of remaining bacteria and bacterial debris.

The strain of *M. smegmatis* used for phage infections was *M. smegmatis* mc^2^155. Mycobacteriophage DyoEdafos was isolated by undergraduate students at the University of Houston Downtown as part of the Science Education Alliance Phage Hunters Advancing Genomics and Evolutionary Science (SEA-PHAGES) Program^44^. Mycobacteriophage ChutneyChips (GenBank PZ440424) was isolated from a soil sample from Mpumalanga, South Africa, using enrichment on a culture of *M. smegmatis* mc^2^155 at 37 °C, and plating for plaques on solid media on a lawn of *M. smegmatis* mc^2^155.

### Phage challenge assays

Phage titer was determined using the small drop plaque assay method^45^. An overnight culture (400 µL) of *B. subtilis* was mixed with 30 mL MMB 0.5% agar supplemented with 1 mM IPTG. Agar was poured into a 10-cm square plate followed by incubation for 1 h at room temperature. Tenfold serial dilutions in MMB were carried out for each of the tested phages and 10 µL drops were put on the bacterial layer. After the drops had dried up, the plates were inverted and incubated overnight at 25 °C. PFUs were determined by counting the derived plaques after overnight incubation and lysate titer was determined by calculating PFUs per mL. When no individual plaques could be identified, a faint lysis zone across the drop area was considered to be 10 plaques. The efficiency of plating was measured by comparing plaque assay results for control bacteria and those for bacteria containing the defense system and/or the defense system with a candidate anti-defense gene.

### Cloning and plasmid construction

Wild-type phage DyoEdafos Adps, Namat, and RyDEP, as well as RyDEP homologs H1–H8 (see Supplementary Table 1 for homolog information), were synthesized (Twist Bioscience) and cloned into custom pET vectors with an N-terminal 6×His-SUMO2-GS tag fusion^46^. To generate mutant DyoEdafos RyDEP constructs, mutations were introduced into the plasmid using site-directed mutagenesis PCR, and the sequences were confirmed by whole-plasmid sequencing (Plasmidsaurus).

### Recombinant protein expression and purification

Recombinant wild-type DyoEdafos Adps, Namat, RyDEP, and RyDEP mutants (Q54A, H75A, H75Q, W78A, H100A, D117A, D117N) as well as the TIR domains of *Aa*TIR (*Aquimarina amphilecti* WP_091411838.1) and *Ab*TIR (*Acinetobacter baumannii* WP_234622687.1) were purified as previously described^46^. In brief, all proteins were cloned into custom pET vector as an N-terminal His-SUMO fusion-tagged protein with a GS linker and transformed into *E. coli* strain BL21-RIL. Large scale cultures (2–4 l) were grown for 5 h at 37 °C (supplementing media with 10 mM nicotinamide), then induced with IPTG overnight at 16 °C. Bacterial pellets were resuspended and sonicated in lysis buffer (20 mM HEPES-KOH pH 7.5, 400 mM NaCl, 30 mM imidazole, 10% glycerol, and 1 mM DTT) and purified using Ni-NTA resin (Qiagen). The Ni-NTA resin was washed with lysis buffer supplemented with 1 M NaCl and then eluted with lysis buffer supplemented with 300 mM imidazole. The Ni-NTA elution fraction was dialyzed into 20 mM HEPES-KOH pH 7.5, 250 mM KCl, 1 mM TCEP overnight while removing the SUMO2 tag with recombinant human SENP2 protease (D364– L589, M497A). All proteins were concentrated using a 10 kDa cut-off concentrator (Millipore) to greater than 10 mg/mL (except *Aa*TIR and *Ab*TIR which were at trace concentrations), flash-frozen with liquid nitrogen, and stored at −80 °C. Proteins used for crystallography were further purified by size-exclusion chromatography using a 16/600 Superdex 75 column (Cytiva) prior to flash-freezing.

### Synthesis of cADPR-based Thoeris signals

Radiolabeled 2′cADPR and 3′cADPR were synthesized using radiolabeled NAD^+^. To generate radiolabeled NAD^+^, synthesis was performed at 37 °C for 20 h in 100 µL reactions containing 20 mM Tris-HCl pH 7.5, 20 mM MgCl_2_, 1 mM nicotinamide mononucleotide, 1 mM ATP, trace amounts of α-^32^P-labeled ATP (Revvity), 1 mM DTT, and 25 µM of the human enzyme NMNAT1 purified from *E. coli* as previously described^47^. Radiolabeled 2′cADPR synthesis was then performed at 37 °C for 20 h in 200 µL reactions containing 40 mM HEPES-KOH pH 7.5, 150 mM NaCl, 50 µM radiolabeled NAD^+^, and trace amounts of the core TIR domain of *Ab*Tir from *Acinetobacter baumannii* purified from *E. coli*. Radiolabeled 3′cADPR was synthesized under the same conditions as 2′cADPR, using trace amounts of the core TIR domain of *Aa*Tir from *Aquimarina amphilecti* purified from E. coli, in place of *Ab*Tir. Following cADPR synthesis, reactions were treated with CIP to remove leftover radiolabeled ATP, boiled for 5 min, and centrifuged for 10 min at 13,000 × g. The supernatant containing the radiolabeled cADPR products was aliquoted and frozen for use as input in biochemical experiments.

### Thin-layer chromatography

cADPR synthesis reactions were used as inputs for downstream degradation reactions, which were carried out as previously described^19^. Briefly, degradation reactions were carried out at 37 °C in 10-µL mixtures composed of 1 µL of a 10× recombinant enzyme stock or cellular lysate, 1 µL of the appropriate synthesis reaction (containing nanomolar concentrations of radiolabeled cADPR signals), 50 mM Tris-HCl pH 7.5, 50 mM MgCl_2_, 2 mM MnCl_2_, 100 mM KCl, and 1 mM DTT. After 20–60 min incubation (when testing purified recombinant enzyme) or 16 h incubation (when testing cell lysates), 0.5 µL volumes of reactions were spotted on a 20 cm × 20 cm PEI cellulose thin-layer chromatography plate (Sigma Aldrich) and developed in 1.5 M KH2PO4 (pH 3.8) buffer for 100 min. Plates were dried at room temperature, exposed to a storage phosphor screen, and detected with a Typhoon Trio Variable Mode Imager System (GE Healthcare).

### EMSA

Electrophoretic mobility shift assay (EMSA) was used to assess 3′cADPR and 2′cADPR immune signal binding activity with DyoEdafos RyDEP wildtype and point mutants, as well as with the eight RyDEP homologs selected for biochemical characterization (H1-H8). Proteins were present at a final concentration of 5 µM in a reaction buffer containing 50 mM Tris-HCl pH 7.5, 50 mM MgCl_2_, 2 mM MnCl_2_, 100 mM KCl, and 1 mM DTT. Proteins were incubated with nanomolar concentrations of α-^32^P-labeled 3′cADPR or 2′cADPR for 1 hour on ice. Reactions were then resolved on a 7.2-cm nondenaturing 6% polyacrylamide gel run at 100 V for 37 min in 0.5× TBE buffer. The gel was fixed in a solution containing 40% ethanol and 10% acetic acid and dried at 80 °C for 1 hour. The dried gel was exposed to a storage phosphor screen overnight and visualization was performed using a Typhoon Trio Variable Mode Imager System (GE Healthcare).

### NAD assay

NAD production was measured using a sequential NARP1 reconstitution assay coupled to a NAD/NADH-Glo biochemical assay (Promega). WT Ry-DEP-cleaved 2′cADPR product, corresponding to 100 µM ADPR-equivalent substrate, was incubated in 50 mM Tris-HCl pH 7.5, 12 mM MgCl_2_, 0.02% BSA, and 0.5 mM ATP. Reactions were assembled with no enzyme, Adps alone, or Adps and Namat, with a matched control using 2′cADPR treated with catalytically inactive RyDEP H75A. For the first reaction phase, samples were incubated with 0.5 µM Adps for 30 min at 30 °C. NAM was then added to all reactions to 0.1 mM, and 0.2 µM Namat was added where indicated for a second 30 min incubation at 30 °C. Completed reactions were diluted 200-fold in reaction buffer and mixed 1:1 with Promega NAD detection reagent in a 96-well plate. After incubation at room temperature, luminescence was measured and NAD concentration was calculated from a standard curve generated with known NAD concentrations.

### Liquid chromatography-mass spectrometry

Reaction conditions for all LC-MS experiments matched those used for TLC analysis, except that unlabeled chemical standards of the corresponding signals were used in place of radiolabeled substrates. Samples were analyzed on an Agilent 1260 HPLC equipped with a diode array detector and an Agilent 6125 single quadrupole mass spectrometer in negative ion mode using a reverse-phase Agilent InfinityLab Poroshell SB-Aq column (2.7-µm particle size, 2.1-mm inner diameter, 150-mm length) at a flow rate of 0.45 mL min^−1^ with a gradient from 100% solvent A (0.1% ammonium formate) to 100% solvent B (methanol). Data were collected by OpenLAB CDS v.2.7 software.

### Cell lysate preparation

*E. coli* and *B. subtilis* phage infected cell lysates were prepared as previously described^19^. In brief, overnight cultures of *E. coli* or *B. subtilis* were diluted 1:100 in 250 mL MMB medium and grown at 37 °C for *E. coli* and 30 °C for *B. subtilis* (250 r.p.m.) until reaching an OD600 of 0.3. The cultures were infected with phages at a final multiplicity of infection of 2. Samples of infected cells were taken before culture collapse. Samples of 5 mL in volume were taken and centrifuged for 5 min at 3,200 × g and 4 °C. The culture pellets were flash frozen using dry ice and ethanol. *E. coli* pellets were resuspended in 250 µL of a lysis buffer containing 20 mM HEPES-KOH pH 7.5, 150 mM NaCl, 5 mM MgCl_2_, 1 mM MnCl_2_, 1 mM DTT, 10% glycerol and 1% NP-40, and incubated at room temperature for 30 min with occasional vortexing. *B. subtilis* pellets were first treated with T4 lysozyme (ThermoFisher) at 1 mg/mL in PBS at 37 °C for 10 min, followed by addition of 400 µL of *E. coli* lysis buffer and 30-min incubation at room temperature. Samples were clarified by centrifugation for 5 min at 17,000 × g at 4 °C, and the supernatant was aliquoted and flash frozen in liquid nitrogen, and stored at −80 °C. For *M. smegmatis* phage infected cell lysates, *M. smegmatis* mc^2^155 was grown to saturation at 37 °C in Middlebrook 7H9 supplemented with 10% albumin dextrose complex (ADC) and 0.05% Tween 80, and for phage infections subcultured into 7H9/ADC supplemented with 1 mM CaCl_2_ and grown at 37 °C with shaking to mid-log phase growth (OD600 ∼0.4). Mycobacteriophage lysates were prepared by picking a well-isolated plaque into 100 µL of phage buffer (10 mM Tris pH 7.5, 10 mM MgSO4, 68 mM NaCl), PCR verified using phage-specific primers, and amplified on solid media. Five mL of *M. smegmatis* culture were infected with 10 µL of phage lysate at a multiplicity of infection of ∼2–3. After incubation at 37 °C for 2.5 hours, cells were collected by centrifugation at 5,000 rpm (∼ 4,300 × g) for 10 minutes at 4 °C, washed with 1 mL of 7H9 medium, and collected by centrifugation at 11,000 × g for 1 minute. The cell pellets were flash-frozen and stored at −80 °C. To lyse the cells, the frozen pellets were thawed, resuspended in 250 µL of a lysis buffer (20 mM HEPES 7.5, 150 mM KCl, 1 mM DTT, 5 mM MgCl_2_, 1 mM MnCl_2_, 10% glycerol) and sonicated intensely 5 times for 15 seconds each with at least 1 minute on ice between rounds. After sonication, the samples were centrifuged at 14,000 × g for 5 min, and the supernatant was obtained, aliquoted, and flash-frozen for storage before use.

### RyDEP Bioinformatic identification

The NARP1 enzymes in DyoEdafos were identified using DefenseFinder^48– 51^. Genome comparison analysis between DyoEdafos and ChutneyChips was performed in Phamerator^52^.

### Identification of RyDEP homologs and generation of phylogenetic trees

Phage DyoEdafos RyDEP belongs to a previously uncharacterized R12-domain protein family designated with the Pfam accession number PF02026^53^. To construct the phylogenetic tree in Figure 4e, viral sequences belonging to the RyDEP Pfam were downloaded and aligned using MAFFT L-INS-i^54^ in Geneious Prime (2025.2.2). The sequences in the region aligned to the R12 domain in RyDEP were extracted and used to construct a phylogenetic tree in Geneious Prime (2025.2.2) using the FastTree plugin. iTOL (v7) was used to annotate the tree^55^.

### Crystallization and structure determination

Crystals of DyoEdafos RyDEP were grown in a hanging-drop format using the hanging-drop vapour diffusion method for 7–9 days at 18 °C. Recombinant DyoEdafos RyDEP was diluted to 10 mg/mL in a buffer containing 500 µM of the substrate 3′cADPR and 20 mM HEPES-KOH pH 7.5, 100 mM KCl, and 1 mM TCEP. The resultant protein mixture was allowed to equilibrate to 18 °C for 10 min and crystals were grown in 96-well trays containing 70 µL reservoir solution and 0.2 µL drops. Drops were mixed 1:1 with purified protein and reservoir solution (0.2 M KSCN, 0.1 M HEPES-NaOH pH 7.5, 20% w/v PEG 3350, and 500 µM 3′cADPR). Crystals were cryo-protected with a reservoir solution supplemented with 22.5% (v/v) glycerol and harvested by flash-freezing in liquid nitrogen.

X-ray diffraction data were collected at Advanced Photon Source Argonne National Laboratory (APS). Data were collected using NECAT-remote GUI v.6.2.3, and processed with XDS, POINTLESS 1.12.16 and AIMLESS 0.7.15^56,57^. Experimental phase information was determined by molecular replacement using Phaser-MR in Phenix and a model of DyoEdafos RyDEP structure generated using AlphaFold3^58,59^. Model building was performed using Coot, and refinement was carried out using Phenix. A summary of crystallographic statistics is provided in Extended Data Table 1. All structural figures were generated using ChimeraX (v1.11.1).

### Nuclear magnetic resonance of RyDEP product

For NMR sample preparation, 2′cADPR standard was reconstituted in water and recombinant RyDEP enzyme was buffer exchanged into water using a 3 kDa concentrator. A large-scale cleavage reaction was assembled at ∼275 µL total volume, containing >10 mM 2′cADPR and ∼300 µM RyDEP enzyme, and incubated overnight at 37 °C. The following day, protein was removed using a 3 kDa concentrator, the flow-through was sampled by LCMS to confirm cleavage, dried completely by SpeedVac, and resuspended in DMSO-d_6_ to yield an approximately 5 mM cleaved product sample for NMR. A Bruker Avance II 600 MHz NMR spectrometer with a cryogenic probe was used to acquire ^1^H, ^13^C, ^1^H-^1^H gCOSY, ^1^H-^1^H NOESY, ^1^H-^13^C HMBC, and ^1^H-^13^C HSQC spectra for the produced ADPR product sample in DMSO-d_6_ at 25 °C.

### Generation of phage DyoEdafos Δ*RyDEP*

DyoEdafos Δ*RyDEP* was constructed using CRISPY-BRED as described previously^60^. In brief, DyoEdafos genomic DNA and a 500 bp dsDNA substrate with the gene 78 (RyDEP gene) deletion were co-electroporated into a recombineering strain of *M. smegmatis* mc^2^155. After 4 hours recovery at 37 °C, cells were then plated onto solid media with kanamycin together with *M. smegmatis* carrying a CRISPR-Cas9 plasmid expressing a sgRNA targeting gene 78. Fifteen primary plaques were picked and screened by PCR, five of which contained the desired deletion. Plaques were purified and sequenced and shown to have the desired loss of gene 78. They also had an unrelated 4,490 bp deletion (coordinates 69,489 to 73,978), which was also present in the parent DyoEdafos, relative to the previously reported sequence (GenBank Accession number MN234187.1).

---

**Extended Data Figure 1.**
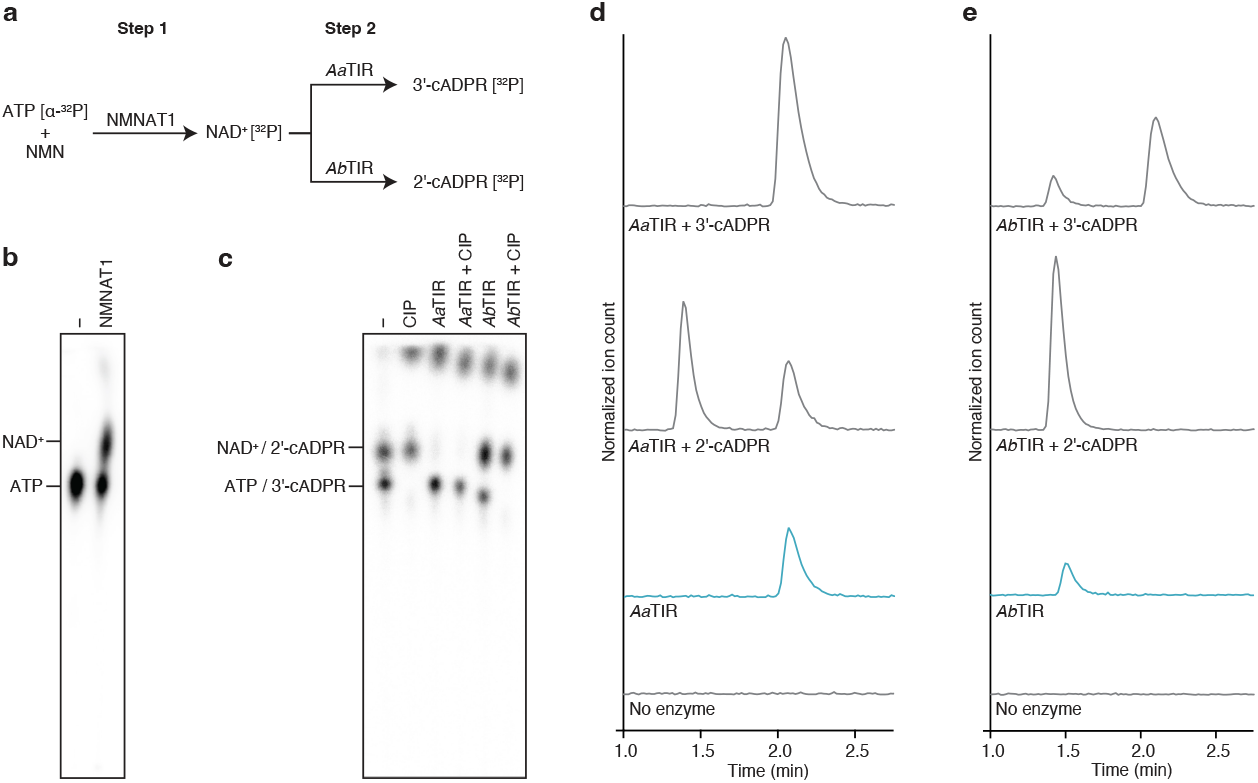
Generation and validation of radiolabeled Thoeris signals. **a**, Schematic depicting the two-step enzymatic reaction developed to generate radiolabeled 2′cADPR and 3′cADPR. **b**, TLC analysis of the first step reaction confirming synthesis of radiolabeled NAD^+^. Data are representative of two independent experiments. **c**, TLC analysis of the second-step reaction confirming synthesis of radiolabeled 2′cADPR and 3′cADPR. CIP treatment degrades unincorporated radiolabeled ATP, ensuring that the primary radiolabeled species present in the final reaction is either 2′cADPR or 3′cADPR. Data are representative of two independent experiments. **d**, LC-MS selected ion monitoring (SIM) traces for cADPR ions (*m/z* 540) showing that the *Aa*TIR reaction product co-elutes with 3′cADPR but not 2′cADPR, consistent with *Aa*TIR 3′cADPR synthesis. Data are representative of two independent experiments. **e**, LC-MS SIM traces for cADPR ions (*m/z* 540) showing that the *Ab*TIR reaction product co-elutes with 2′cADPR but not 3′cADPR, consistent with *Ab*TIR 2′cADPR synthesis^6,7^. Data are representative of two independent experiments.

**Extended Data Figure 2.**
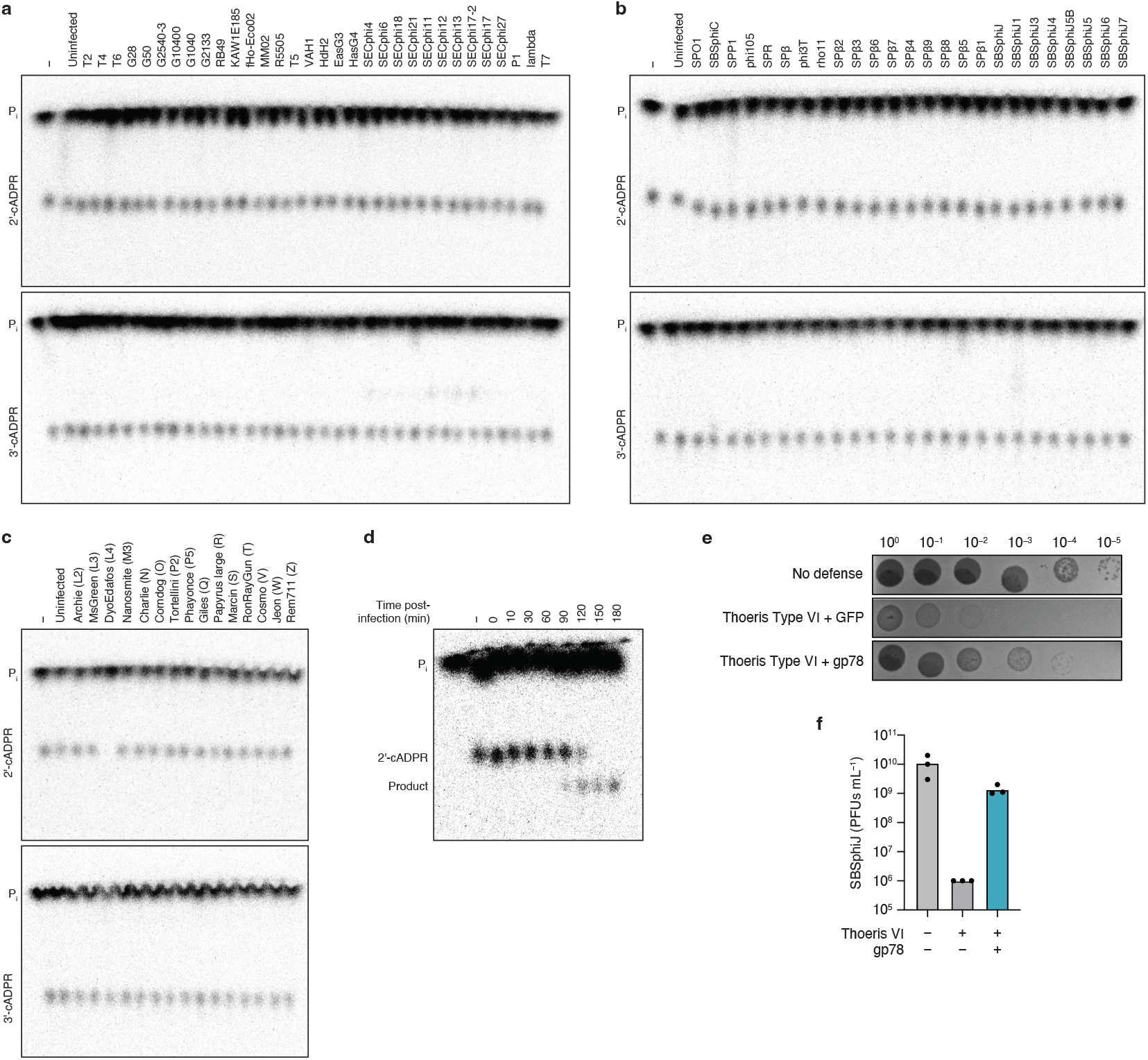
A biochemical screen reveals that DyoEdafos phage encodes for a Thoeris immune evasion enzyme. **a, b, c**, TLC screen for 2′cADPR/3′cADPR degrading activity in *E. coli* (**a**), *B. subtilis* (**b**), and *M. smegmatis* (**c**) phage infected lysates. Data are representative of two independent experiments. **d**, TLC analysis of 2′cADPR degradation by phage DyoEdafos-infected lysates harvested at different times post-infection. Reduced mobility of the final product is likely due to additional processing that occurs in complex lysates following RyDEP-mediated cleavage. Data are representative of two independent experiments. **e, f**, Plaque assay (**e**) and quantification (**f**) of *B. subtilis* co-expressing a Type VI Thoeris operon that signals through 2′cADPR with DyoEdafos RyDEP or a GFP control against serial dilutions of phage SBSphiJ. Data represent three independent experiments.

**Extended Data Figure 3.**
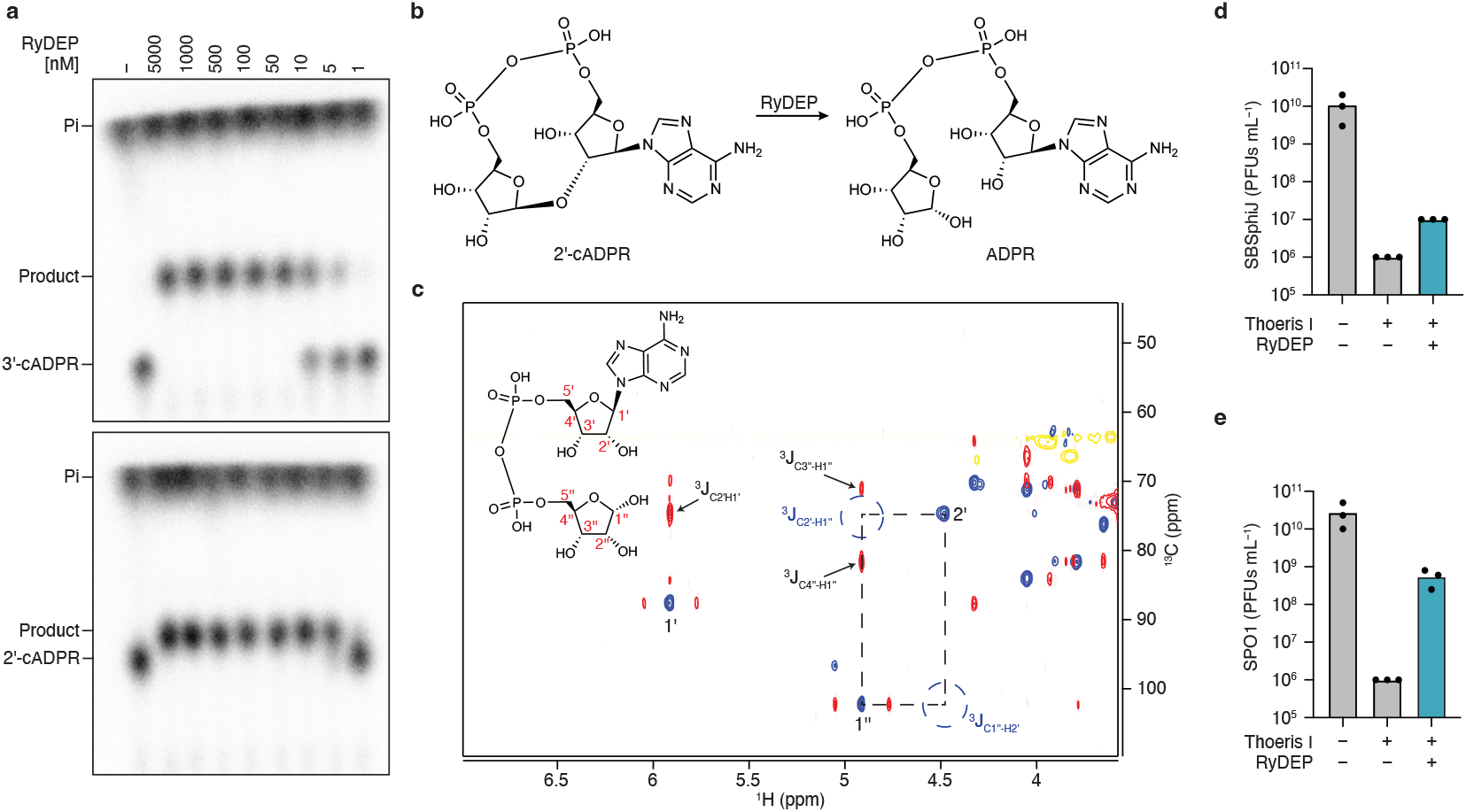
RyDEP is a 2′cADPR-preferring glycosidase that broadly counteracts cADPR-mediated Thoeris defense. **a**, TLC analysis of RyDEP titration reactions against 1 µM of 2′cADPR or 3′cADPR. Reactions were incubated for 20 min and stopped by boiling. Data are representative of two independent experiments. **b**, Schematic of reaction performed by RyDEP. **c**, ^13^C-HMBC spectrum of the RyDEP cleavage product, with HSQC peaks overlaid in blue. The absence of HMBC correlations corresponding to the intact ribose–ribose linkage (as indicated by dashed blue circles) supports cleavage of the glycosidic bond, while the observed HMBC peak pattern (as indicated by the black markers) is consistent with a β-anomeric adenosine ribose and an α-anomeric terminal ribose. **d, e**, Quantification of phage SBSphiJ (**d**) or phage SPO1 (**e**) replication in *B. subtilis* co-expressing a Type I Thoeris operon that signals through 3′cADPR with DyoEdafos RyDEP or a GFP control. RyDEP-mediated evasion is comparable for phage SPO1 in type I and type VI Thoeris, but stronger for phage SBSphiJ against type VI than type I, consistent with RyDEP preference for 2′cADPR. Data represent three independent experiments.

**Extended Data Figure 4.**
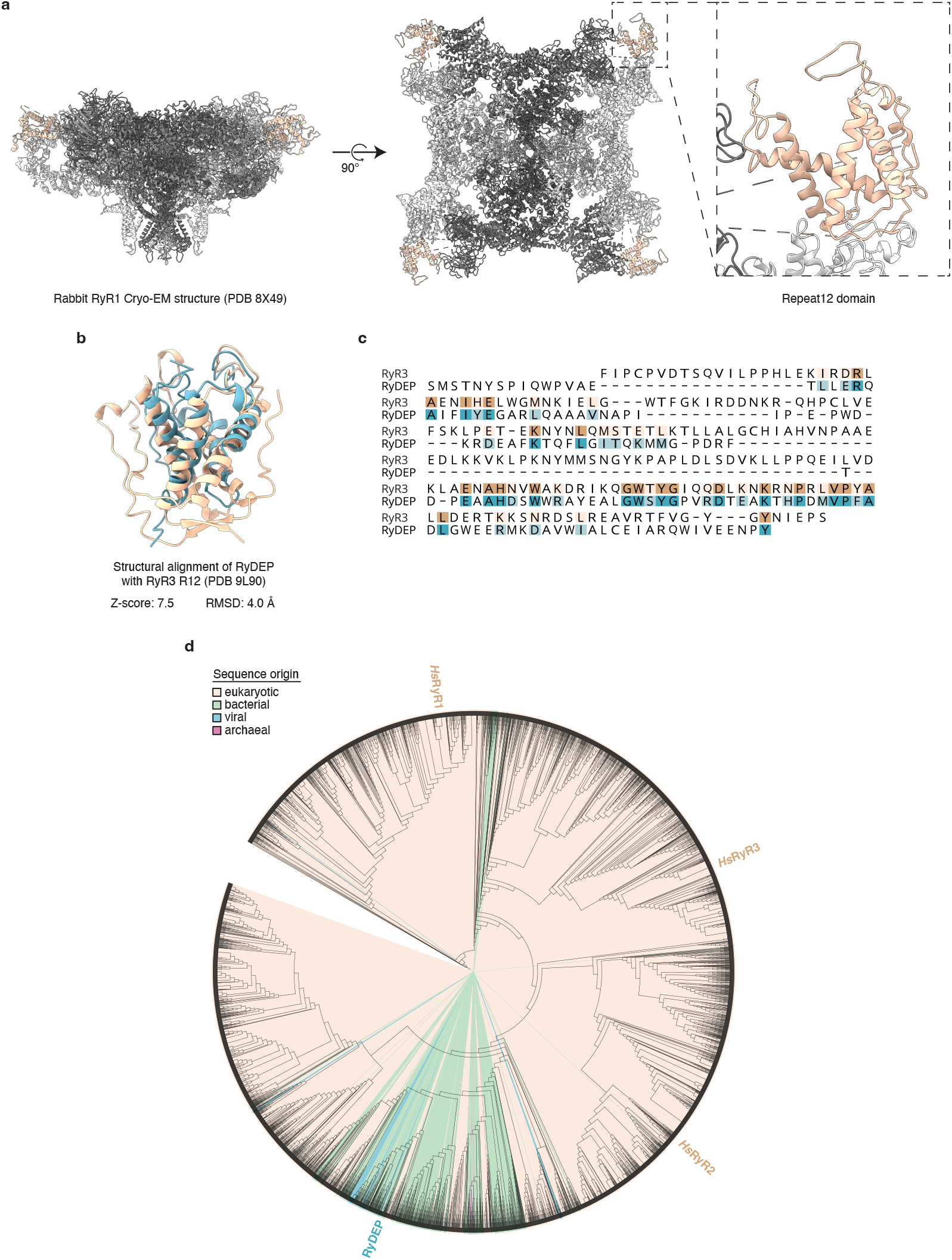
RyDEP is homologous to the R12 domain of animal RyRs. **a**, Overall structure of an animal RyR homotetramer, with a zoomed-in view of the R12 repeat located at the corners of the complex. **b**, Structural superposition of phage DyoEdafos RyDEP and the R12 repeat from human RyR3 generated using the RCSB PDB pairwise structure alignment tool. **c**, Structure-derived residue alignment from the same comparison, with structurally aligned residues highlighted. **d**, Phylogenetic tree of R12-domain proteins (Pfam PF02026) colored by sequence origin, with RyDEP and human RyRs indicated at tree nodes. Proteins were aligned using MAFFT FFT-NS-1 in Geneious Prime (2025.2.2), sequences in the region corresponding to the RyDEP R12 domain were extracted, and the tree was built using the FastTree plugin in Geneious Prime (2025.2.2) and annotated in iTOL.

**Extended Data Figure 5.**
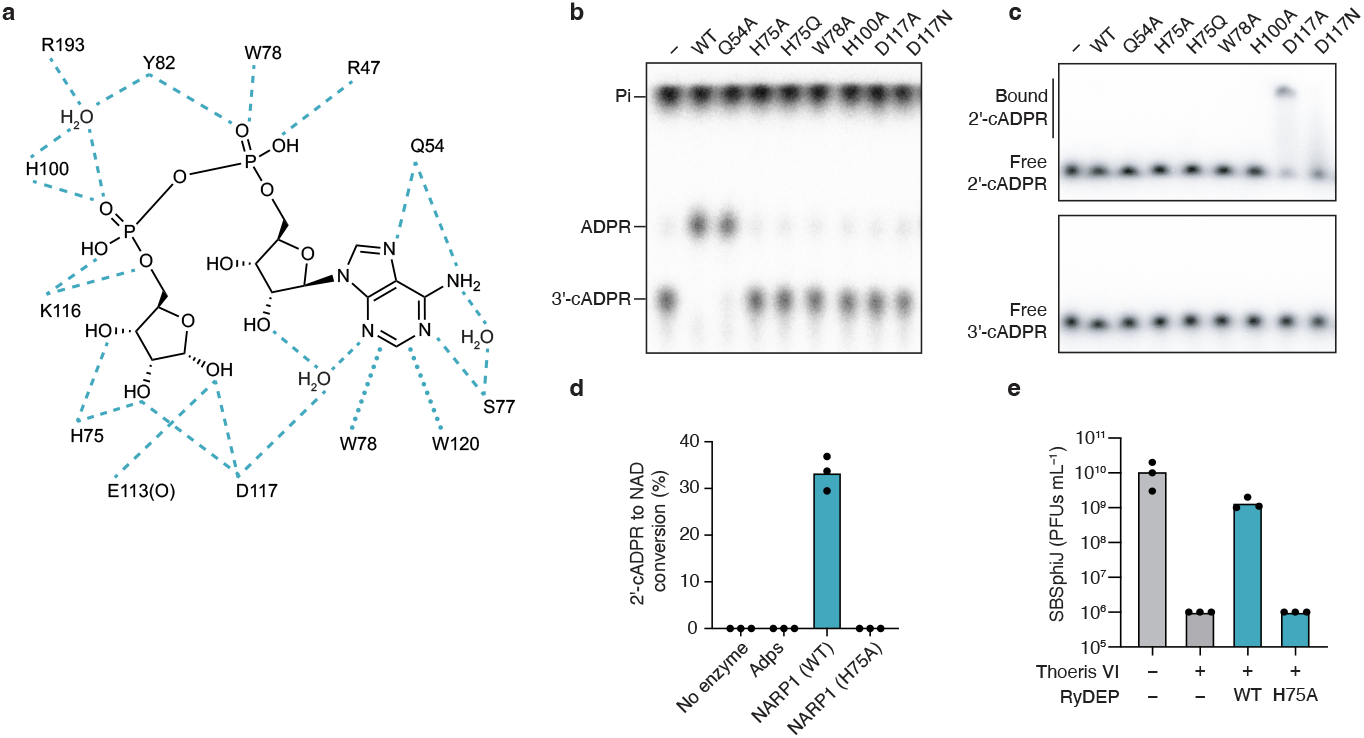
Structural basis of 3′cADPR hydrolysis by RyDEP. **a**, Schematic depicting the polar contacts (dashed lines) and non-polar contacts (dotted lines) between RyDEP and linear ADPR in the post-reaction state. **b**, TLC analysis of the catalytic activity of RyDEP point mutants with 3′cADPR. Data are representative of two independent experiments. **c**, EMSA analysis of the binding ability of RyDEP point mutants to 2′cADPR or 3′cADPR. Data are representative of two independent experiments. **d**, Conversion of 2′cADPR into NAD+ by purified phage DyoEdafos NARP1 enzymes and WT RyDEP or the catalytically inactive H75A mutant RyDEP. NAD+ concentrations were determined using the NAD/NADH-Glo assay. Data represent three independent experiments. **e**, Plaque assay quantification of phage SBSphiJ replication in *B. subtilis* co-expressing Type VI Thoeris with wild-type RyDEP or the catalytically inactive H75A mutant. Data represent three independent experiments.

**Extended Data Figure 6.**
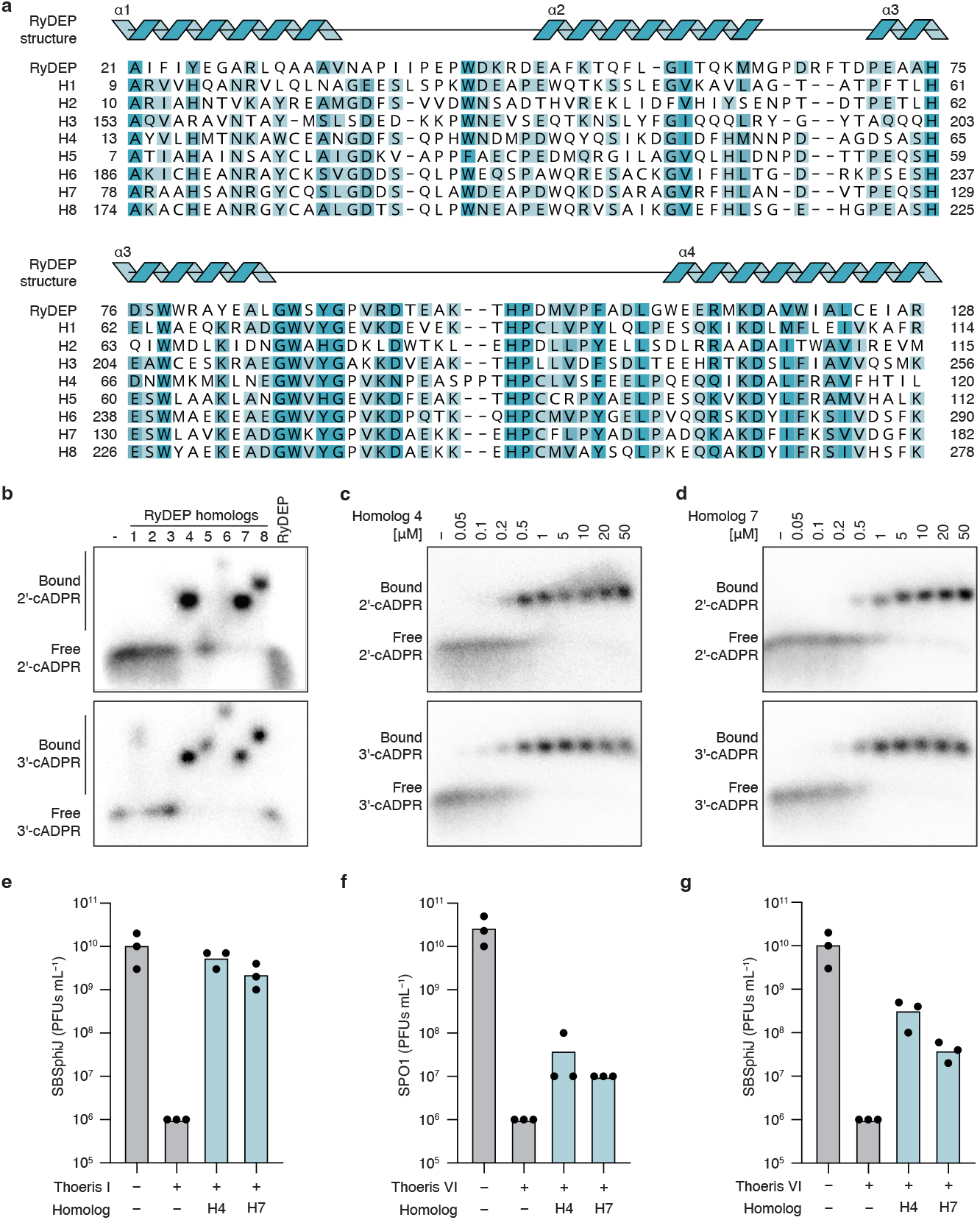
Phages encode non-enzymatic RyDEP homologs that function as anti-Thoeris sponges. **a**, Sequence alignment of all eight homologs purified for biochemical characterization, with conserved residues highlighted in blue. The RyDEP secondary structure is mapped above the alignment, with helices shown as blue cartoons. **b**, EMSA binding analysis showing the ability of each RyDEP homolog to form complexes with 2′cADPR and 3′cADPR (summarized in Fig. 4f). Data are representative of two independent experiments. **c, d**, EMSA analysis of H4 (**c**) and H7 (**d**) binding to 2′cADPR and 3′cADPR. Data are representative of two independent experiments **e**, Plaque assay quantification of phage SBSphiJ replication in *B. subtilis* co-expressing Type I Thoeris with the selected RyDEP homologs H4 and H7 or a GFP control. Data represent three independent experiments. **f, g**, Plaque assay quantification of phage SPO1 (**f**) or phage SBSphiJ (**g**) replication in B. subtilis co-expressing Type VI Thoeris with the selected RyDEP homologs H4 and H7 or a GFP control. Data represent three independent experiments.

