## Extended Data Table 1 for "Phage RyR-domain proteins degrade ADPR-based immune signals and fuel NAD^+^ synthesis"

**Table 1 Data collection and refinement statistics (molecular replacement)**

| | DyoEdafos RyDEP–3′cADPR post reaction state |
|---|---|
| **Data collection** | |
| Space group | P $2_1$ $2_1$ $2_1$ |
| Cell dimensions | |
| *a*, *b*, *c* (Å) | 60.825 67.199 68.297 |
| α, β, γ (°) | 90 90 90 |
| Resolution (Å) | 68.30 – 1.65 (1.68 – 1.65) * |
| $R_{merge}$ | 0.197 (1.679) |
| $I / \sigma I$ | 8.0 (1.5) |
| Completeness (%) | 100.0 (100.0) |
| Redundancy | 13.1 (12.4) |
| **Refinement** | |
| Resolution (Å) | 47.90 – 1.65 |
| No. reflections | 34244 |
| $R_{work}$ / $R_{free}$ | 0.1836 / 0.2195 |
| No. atoms | |
| Protein | 2256 |
| Ligand/ion | 72 |
| Water | 358 |
| *B*-factors | |
| Protein | 19.92 |
| Ligand/ion | 17.19 |
| Water | 30.93 |
| R.m.s. deviations | |
| Bond lengths (Å) | 0.014 |
| Bond angles (°) | 1.15 |

*Values in parentheses are for highest-resolution shell. Data were collected from a single crystal.
